# Temporal Dynamics of Urinary Extracellular Vesicle Excretion and Cargo in Healthy Subjects over 24 Hours

**DOI:** 10.1101/2025.06.17.660244

**Authors:** S Upson, M Selesky, M Greig, H Yavuz, M Harding, I Mallawaarachchi, J Ma, M Solga, A Alwagdani, L Musante, U Erdbrügger

## Abstract

Urinary extracellular vesicles (uEVs) are promising non-invasive biomarkers of renal physiology and pathology, yet little is known about their temporal excretion patterns or optimal normalization strategies as urine concentrations vary significantly during the day. In this basic physiology study, we used a uniquely designed processing protocol of urine collection to characterize the variation of uEV excretion and cargo throughout a 24 hour period in healthy subjects. Thirteen healthy adults collected every individual void over 24 hours and these voids were independently analyzed along with a composite sample intended to recreate a 24 hour collection sample. uEVs were enriched by low-speed centrifugation with three low-ionic-strength washes to minimize uromodulin contamination. Size, concentration, morphology and cargo of uEVs were characterized with nanoparticle tracking analysis, cryo-electron microscopy, immunoblotting and single EV flow cytometry. We observed a stable urine creatinine excretion into the urine over the day. uEV concentrations varied up to 100-fold across the day between individuals with a decreasing trend over time with the result close to reaching statistical significance (p=0.06). uEV concentration was positively associated with urine creatinine, specific gravity, and TSG101, an EV marker. The association between urine flow and uEV excretion varied across the day. Most tested uEV cargo markers exhibited substantial within individual variation over the day without consistent trends in our healthy cohort. When grouped into predefined time periods to visualize temporal patterns, a transient dip at noon was observed, followed by a significant increase in the subsequent period, consistent with a pattern previously reported. These findings offer foundational insights into uEV excretion patterns over the course of a day in a healthy cohort and emphasize key methodological considerations essential for advancing uEV biomarker research.

## Introduction

Urinary extracellular vesicles (uEVs) are a heterogeneous group of membrane-surrounded-vesicles which originate mostly from kidney cells but also from other parts of the urogenital tract^1^. As they carry markers and bioactive molecules of their parent or cell of origin, they are seen as potential biomarkers and cell-to-cell messengers in health and disease^1^. Yet, there has been minimal investigation of how uEV excretion and release into the urine varies throughout the circadian cycle in healthy individuals. Circadian rhythms have been shown to control many physiologic processes in the human body. The kidney has its own circadian clock that works in synergy with the body to maintain homeostasis^32^. About 20% of the genes expressed within the kidney are regulated in a circadian manner, therefore variations in biomarker excretion over the day needs to be considered^33^. However, the circadian rhythm of only a few urinary biomarkers, EV and non-EV related, have been well characterized^36^. The field of translational uEV investigation is impacted but also complicated by the fact that urine is the most dynamic biofluid, and the field currently lacks standardized normalization protocols^35^. Urine contents vary in composition between and within individuals, not only from day to day but also hour to hour. Therefore, these physiologic variations in uEV excretion and normalization methods must be understood to explore the full diagnostic potential of uEVs.

Since a landmark study published in 1983 in the New England Journal of Medicine (NEJM) demonstrated that the calculated protein-to-creatinine ratio of the second-morning void provided an accurate quantification of daily proteinuria, clinicians have been able to use a spot urine collection to extrapolate kidney health and function across a greater span of time than that during which the single void was produced^9^. This approach relies on the fact that the urinary creatinine excretion rate is constant within an individual over time and across individuals, and that the biomarker production or excretion has a linear relationship with urinary creatinine across individuals^34^. This method can therefore not be used for example in acute kidney injury when creatinine excretion is changing from hour to hour.

In this basic physiology study, we used a uniquely designed collection protocol of urine to characterize with great detail the variation of uEV excretion and uEV cargo throughout a 24 hour period in healthy subjects. Thirteen healthy subjects separately collected each spontaneous void throughout a 24 hour period and documented the time of their voids along with a food and drink log for the same time period. Knowledge of the exact time periods of the urine collections allows to test foremost if urinary creatinine excretion is stable or dynamic in our healthy cohort as well as the trends of uEV excretion and uEV cargo over the day. In addition, this study separated and characterized uEVs’ size, morphology and cargo over the course of a 24 hour period to identify temporal patterns of uEV excretion and develop strategies to normalize uEV concentration and cargo in urine which is one of the most dynamic biofluids. Finally, yet importantly, our results were put in context of some of the subject’s food intake.

## Methods

### Study cohort

Healthy subjects above 18 years of age were recruited through advertisement flyers and by word of mouth. All subjects had no history of diabetes mellitus, hypertension, or chronic kidney disease. Subjects were on an unrestricted diet, and nine recorded their dietary intake for the 24 hour collection period using the Automated Self-Administered 24-Hour Dietary Assessment Tool^3^ (ASA24). ASA24 output included kilocalories (kcal), protein, total fat, carbohydrates, and water, as well as vitamins and minerals. The study protocol was approved by the University of Virginia’s Institutional Review Board (IRB#220053). All subjects provided written and verbal informed consent prior to participating in the current study.

### Urine collection and storage

Healthy subjects were given an insulated bag with 5-6 ice packs and urine collection bottles. Subjects kept the ice packs in the freezer and added them to the bag once they began collecting individual voids. These ice packs kept the insulated bags at a temperature of 4°Celsius. Every void was collected during a 24 hour period for each healthy subject, starting with the second void after awakening and ending with the first void of the following morning. Each full void was collected in a separate bottle and labeled with the time of void initiation. After the final void (the first void of the following day), the finished collection was delivered to our laboratory and processed in the lab within two to six hours of the final void. Twenty percent of each void was combined to produce an artificial 24 hour collection sample. In 7 out of 13 patients, serum creatinine level was also obtained on the day urine collection finished.

### Chemical analysis of urine

After the volume of each sample was measured and an aliquot of 20% removed from each void, a urinalysis dipstick (Siemens Multistix 10 SG) analysis was performed in each raw urine sample to assess pH, glucose, bilirubin, ketone, specific gravity, blood, protein, urobilinogen, nitrite, and leukocytes. All data was securely stored in a REDCap database.

Urine protein and creatinine concentrations were also measured in the raw urine aliquots from each void. Protein quantification was performed by Coomassie micro assays on a 96-well plate. The protein in the urine binds to Coomassie Brilliant Blue G-250 which is shown with an absorption at 595 nm^4^. Standard controls were loaded in triplicate on the plate to construct a standard curve for quantification of urine protein content. Plates were scanned at 595 nm to determine absorbance.

Creatinine concentrations were measured utilizing a picrate alkaline Jaffe assay on a 96-well plate^5^. Creatinine and picric acid form a complex that has an absorbance at about 520 nm. Standard controls were loaded on the first three lanes of the plate to make a standard curve to compare urine creatinine content to. Plates were scanned at 520 nm to determine absorbance.

### uEV separation

uEVs were enriched from each individual urine sample, including the artificial 24-hour collection samples, by differential centrifugation. 42 mL of raw urine was centrifuged at a Relative Centrifugal Force (RCF) of 4,600 g at max radius 168 mm (5000 rpm) in a TX-400 Sorvall ST16R (Thermo Fisher Scientific) swing bucket rotor (k Factor 9153) for 30 minutes at room temperature (RT). The supernatant 4,600 g (SN4,600) was then centrifuged in a Sorval SS-34 rotor in a Sorvall RC 5B centrifuge for 30 minutes 16000 rpm at 4 degrees. Then the supernatant was discarded and the pellet was resuspended in a low ionic strength buffer (10 mM HEPES pH 7.4) supplemented with EDTA (2.5 mM pH 8.0) to remove uromodulin and transferred into an Eppendorf tube to be spun at 15,200 RPM for 30 minutes at room temperature in an Eppendorf microcentrifuge fixed angle rotor (Thermo Scientific, 75003602) in a Sorvall ST16R centrifuge (Thermo Scientific) at 4 degrees to generate an EV pellet called P21 as described in Musante et al.^30^ (**Fig. 1**). Two more washes were done with the low ionic strength buffer for a total of three washes. This method allows to generate a relatively pure uEV fraction.^6^ Each of the four pellets generated were equivalent to 10.5 mLs of raw urine to perform characterization and cargo analysis on. 1 mL of raw urine was frozen down for protein and creatinine assays.

**Figure 1:**
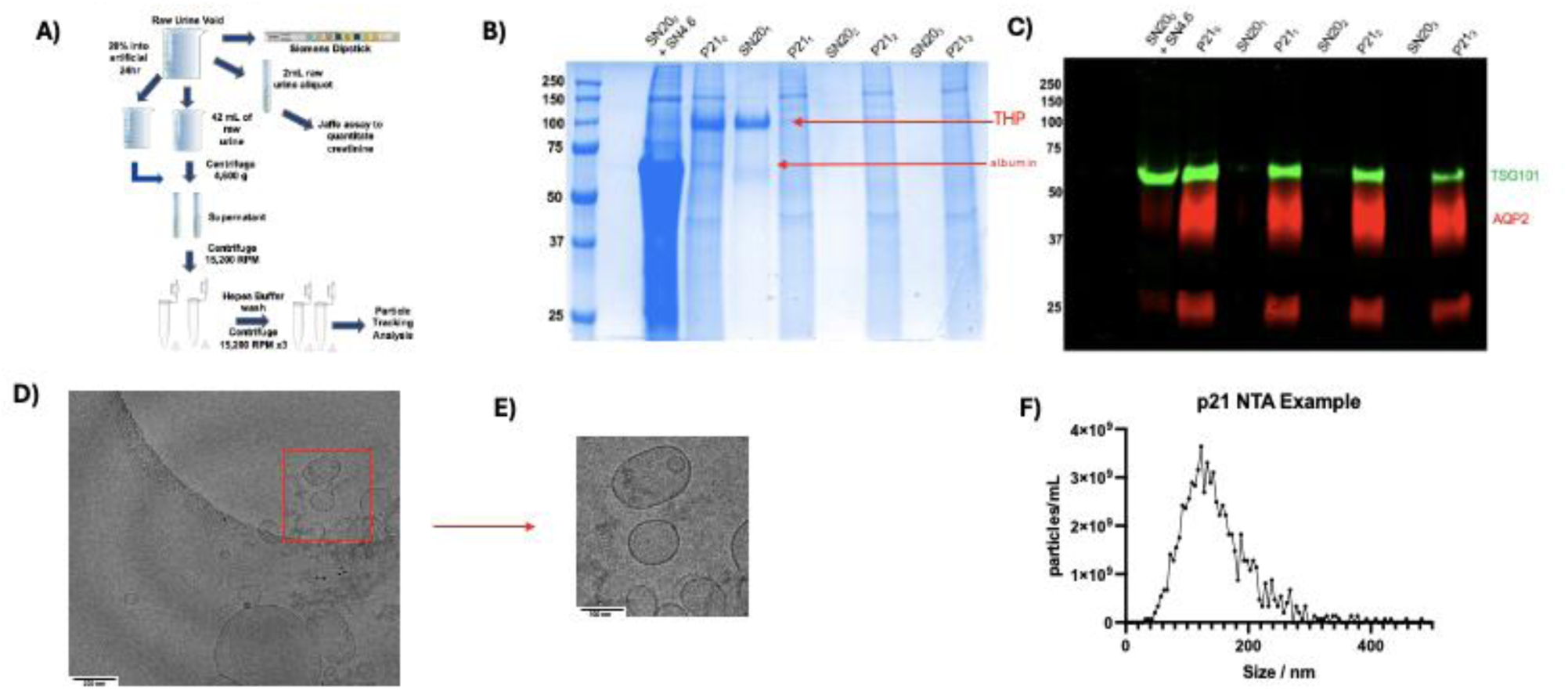
Panel A is a flow of the urine processing methods. Verification of THP reduction in p20 prep. Panel B was stained with Coomassie biue to visuslize proten content. After being spun in a polycarbonate tube at 15,200 RPM for 30 min, the pellet was resuspended in SN4.6 to transfer into an Eppendorf tube to then spin again at 15,200 RPM to visualize the washing steps which resulted in SN20_0_ + SN_4.6_. After one wash with the low ionic srength bufler, the THP is released into the supernatant SN20_1_, Pane C was stained wit direct conjugate AQP2 680 RD, prmary rabbit TSG101, anti rabbit 800 CW. Panel D is cryo electron microscopy of the p20 pellet showing a clean prep. Panel E is a zoom in of D. Panel F shows an example of the p21 pellet size distribution on NTA.

### uEV physical characterization

Urinary EVs were further characterized with a Nanoparticle Tracking Analysis (NTA) tool to assess size and concentration of uEVs in the uromodulin depleted uEV pellets. Cryogenic transmission electron microscopy (Cryo-TEM) was performed to assess uEV morphology and purity of EV prep, ruling out other protein contamination of the uEV prep.

NTA was performed using the ZetaViewPMX 120 (Particle Metrix) configured with a 488 nm laser with a long wave-pass (LWP) cut-off filter (500 nm) and a sensitive CMOS camera 640 x 480 pixels. Each P21 sample was diluted in 2 mL of 0.1 μm filtered (Minisart high flow hydrophilic 0.1 μm syringe filter Sartorious) deionized water (DI 18 MΩ/cm) to obtain a particle concentration between 1 x 10^7^ and 1 x 10^8^ particles/mL. The instrument was set to a constant temperature of 25°C, a sensitivity of 80, a shutter speed of 100, and a frame rate of 30 frames per second (fps). Each sample was measured at 11 different positions throughout the cell, with 5-7 cycles of readings at each position in order to have a minimum of 1000 traces. Post-acquisition parameters were set to a minimum brightness of 30, a maximum size of 200 pixels, and a minimum size of 5 pixels. Automated cell quality control was checked using high-quality deionized water (DI). Camera alignment and focus optimization were performed using polystyrene 100 nm beads (Applied Microspheres). Data analysis was performed with ZetaView 8.05.12 software provided by the manufacturer. Automated reports of the particles recording across the 11 positions were manually checked, and any outlier position was removed to calculate particle concentration and distribution expressed by mode, median, and mean (**Supplemental Figure 1**).

uEV morphology was assessed using Cryo-TEM at the molecular electron microscopy core at the University of Virginia. Low-speed centrifuged uEV P21 pellets were solubilized in 20 μL PBS–0.1 μm and applied to a glow-discharged, perforated carbon-coated grid (2/2-3C C-Flat; Protochips). Low-dose images were collected at a nominal magnification of 29,000 × on the Tecnai F20 Twin transmission electron microscope operating at 120 kV. Digital micrographs were recorded on a Gatan US4000 charge-coupled device camera.

### uEV cargo analysis

Urinary EV cargo was assessed utilizing immunoblotting and single EV flow cytometry following the guidelines by the International Society for Extracellular Vesicles (ISEV)^7,8^. For immunoblotting, P21 pellets equivalent to 10.5 mL of raw urine were solubilized in an electrophoresis solubilization buffer (ESB): 6 M urea, 2 M thiourea, 5% (w/v) sodium dodecyl sulphate (SDS), 40 mM Tris-HCl, pH 6.8, 0.5 mM ethylenediaminetetraacetic acid (EDTA), 20% (v/v) glycerol, supplemented with 50 mM DTT (dithiothreitol) and protease and phosphatase inhibitor cocktail. Samples were run on 4–15% gradient polyacrylamide gels (Bio-Rad, 5671084) according to the manufacturer’s directions. A molecular weight marker ladder was run beside the samples. Protein on the gels was transferred to PVDF membranes for subsequent staining for total protein content (REVERT total protein stain from Li-Cor), and then sequentially probed with antibodies to NCC, aquaporin-2 (AQP2), and TSG101. Imaging and quantification of signal strength of blots were performed on a Li-Cor Odyssey CLx instrument using Image Studio software.

Spectral flow cytometry was performed using the Cytek Aurora 5-Laser flow cytometer equipped with the Enhanced Small Particle (ESP™) Detection tool. P21 samples were prepared from 1 mL of thawed sample using differential centrifugation (4600xg for 30 minutes, preceding 21,100xg for 30 minutes). Data acquisition was standardized by time, with events collected over a 2-minute interval. uEVs were characterized using spectral flow cytometry and analyzed with FSCExpress software to determine total particle counts based on surface marker expression. uEV concentrations were quantified using the instrument’s built-in volumetric method. Specific markers were used to identify uEVs originating from distinct nephron segments based on single gene analysis, including CD26/dipeptidyl peptidase 4(DPP4)(proximal tubular), CD35/complement receptor 1(CR1) (podocyte), CD10/MME/neprilysin (proximal tubular S1, S2, and S3, podocyte), and tetraspanin, CD9 (collecting duct intercalated cell and collecting duct principal). Location determined by single cell expression^37^. Particle sizing was performed using FCMpass software, which estimates particle size using NIST traceable polystyrene particles of known diameters and refractive indices. Scatter intensity data were calibrated into standardized units of scattering cross-section (nm^2^) and particle diameter (nm), enabling phenotypic analysis of medium-sized (mean size: 233.15 nm) and large (mean size: 973.3 nm) EV populations, based on software-generated size distribution curves. Final size distributions were generated by extrapolating diameters from raw SSC data.

## Statistical Analysis

All graphs were generated using GraphPad Prism 10 (version 10.0.0, GraphPad Software). Since each healthy subject contributed multiple voids over the day, statistical analyses accounted for within-subject correlations. Relationships between two quantitative variables and individual biomarkers were evaluated using linear mixed-effects models in R (version 4.5.0), implemented with the lme4 package (version 1.1-37), with Individual included as a random effect.

### Four-Model Comparisons for Relationships Between Two Quantitative Variables

For each pair of variables (e.g., creatinine and uEV concentration, specific gravity and uEV concentration, creatinine and TSG101 signal, uEV excretion and TSG101 signal, urine production and uEV excretion), four candidate models were compared. Models sequentially included fixed effects of the primary predictor for that variable pair, Hour, and the interaction of the predictor and Hour, with random intercepts and random slopes for Individual. Model selection was based on fixed-effect significance, random-effect structure, and convergence. In most cases, the interaction and random slope terms were not significant, or variance was near zero, so final models included only the significant fixed effects (which were often the primary predictor) and a random intercept for Individual

### Two-Model Comparisons for One Quantitative over time

For each biomarker, Median diameter, uEV concentration, uEV concentration (creatinine-normalized), uEV excretion, NCC signal, NCC/TSG101, AQP2 signal, AQP2/TSG101, CD10 count/mL, CD10/creatinine, CD9 count/mL, CD9/creatinine, CD35 count/mL, CD35/creatinine, CD26 count/mL, and CD26/creatinine, two mixed models were fit: (1) Hour as a fixed effect with a random intercept for Individual and (2) the same model including a random slope for Hour. Random slope models were assessed for singularity and whether the slope variance was meaningfully greater than zero. The final models included only a fixed effect of Hour and a random intercept for Individual.

### Nonparametric Analyses of Biomarker

To evaluate within-day variability, nonparametric one-way ANOVA (Friedman tests) assessed protein signals, uEV size, and uEV concentration across five periods: Period A, second morning void; Period B, all voids between Period A and 12 PM; Period C, 12 PM to 6 PM; Period D, 6 PM to 12 AM; and Period E, 12 AM to the next Period A. This approach standardizes time-of-day comparisons across individuals with the periods adapted from Ginsberg et al. Friedman tests were conducted in Prism. Differences between individual void or period values and the 24-hour pooled sample were assessed using Wilcoxon rank sum tests.

## Results

### Study cohort

Thirteen healthy subjects were successfully recruited for this study. These 13 healthy subjects (9 F, 4M) had an average age of 44.3 +/- 11.7 years, BMI of 21.7 +/- 1.79 kg/m^2^, and collected an average of 7.15 +/- 1.57 voids with an overall urine volume of 2010 +/- 734 mL over 24 hour (**Table 1**). For the subset of healthy subjects who got their serum creatinine measured at the time of collection, the estimated glomerular filtration rate was between 87 and 116 ml/min/1.73m^2^ using the CKD-EPI creatinine equation^10^ whereas the calculated creatinine clearance by 24 hour urine collection was between 77.8 and 133.8 mL/min^11^. All clinical characteristics are within the normal range as expected. Clinical characteristics for each of the healthy subjects are included in **Supplemental Figure 2.**

**Table 1:** Overview of 13 healthy individuals’ clinical characteristics and a subset of 7 individuals with serum creatinine values.

| Sex (F, M) | Age | BMI | Body Surface Area | Number of Voids | Urine Flow | Urine Volume |
| --- | --- | --- | --- | --- | --- | --- |
| 9,4 | 44.3 +/- 11.7 yrs | 21.7 +/- 1.79 kg/m<sup>2</sup> | 1.79 +/- 0.163 m<sup>2</sup> | 7.15 +/- 1.57 voids | 0.805 +/- 0.375 mL/min | 2010 +/- 734 mL over 24 hours |
| Subset of 7 individuals |  |  |  |  |  |  |
| Serum Creatinine |  |  | calculated GFR by 24 hr collection |  | estimated GFR by CKD-EPI |  |
| 0.6 – 0.9 mg/dL |  |  | 93-159 (ml/min/1.73m<sup>2</sup>) |  | 87-116 (ml/min/1.73m<sup>2</sup>) |  |

### EV Characterization of the p20 pellet

To reduce contamination with uromodulin, the most abundant urinary protein in healthy urine, the uEV pellets were generated with a low centrifugation spin and washed with low ionic strength buffer three times. Figure 1E shows a successful reduction of uromodulin without compromising the EV quantity during the process of washing. The reduction of uromodulin filaments with the three washes of low ionic strength buffer is demonstrated in the Coomassie Gel as well as the preservation of TSG101 signal. The final pellet P21_3_ (after 3 washes) is clean with the reduction of uromodulin at 100 kd (**Fig. 1B**). The EV marker TSG101 and AQP2 signal is minimally diminished after the washing steps **(Fig. 1C**). Cryo-TEM shows uEV p21 pellet without protein filaments “entrapping” the vesicles as well as the heterogeneity of the size and shape of uEVs (**Fig. 1D**). The final product of our protocol utilizing three washes is a “cleaned” uEV prep which should lead to a significant reduction in non-EV particles. Therefore counts of particles detected per NTA are very close to the actual EV counts. An example of the size distribution of uEV sample is shown in **Fig. 1F**.

### uEV concentration, excretion and size variation over 24 hour period

Nanoparticle Tracking Analysis (NTA) was utilized to count and size uEVs. As described above, our uEV separation protocol generates uEV pellets free from contamination with other protein complexes, as verified with cryo electron microscopy (Fig. 1). We therefore exchanged the term particles with uEVs in our summary: uEV concentration varies over the course of a day from 6.2 * 10^6^ to 4.5 * 10^9^ particles/mL with a slight decreasing trend uEV concentration over time (**Fig. 2A**,β = –0.018, p = 0.0601), but this did not reach statistical significance. uEV concentration normalized to creatinine varies over the course of a day from 1.1 * 10^10^ to 1.6 * 10^12^ particles/gram of creatinine with again a slight decreasing trend uEV concentration over time (**Fig. 2B**, β = – 0.0136, p = 0.066). When voids were grouped into the prespecified time periods adapted from Ginsberg et al, the difference in uEV concentration was up to 100-fold, though the changes were not statistically significantly different (**Fig 2C**, p=0.622). Individual voids (1-7) of uEV concentration were also compared to the 24 hr sample to test for differences but no comparisons were significantly different (**Supplemental Fig 3**). uEV excretion rate over the day varies from 2.4 * 10^6^ to 2.2 * 10^9^ particles/minute but the decreasing linear trend was weak not reaching significance(**Fig 2D)**. However, additional analysis was performed to investigate if excretion varied during morning or night (**Supplementary Fig 5**). A linear mixed model revealed significant effects of urine production (β = 0.42, p = 0.0003) and time of day (β = 0.043, p = 0.018) on uEV excretion rate. Importantly, the urine production (ml/min) × hour interaction was significant (β = –0.0365, p = 0.00016), indicating that the association between urine flow and uEV excretion varied across the day (**Supplemental Fig 5F**). Predicted values show that urine flow increases uEV excretion in the early morning, with a progressively weaker effect at midday and decreasing in evening (**Supplement Fig 5E**). Thus, both urine production and time of voiding jointly contribute uEV excretion dynamics. TSG101 signal was strongly positively associated with urinary EV excretion rate (**Fig 2E** β = 1.10, p = 1.2×10⁻⁶), indicating that TSG101 levels scale proportionally with the number of EVs excreted. NTA demonstrated median uEV size ranged between 93.9 to 220.1 nm, but median size showed no association with time of day (**Fig 2F** β = 0.001, p = 0.997), indicating that median size remained stable throughout the day with no trend. With Period grouping, again differences across a 24 hour period were not significant (p=0.150, **Fig. 2G**). Individual voids (1-7) of uEV median size were also compared to the 24 hr sample to test for differences but no comparisons were significantly different (**Supplemental Fig 4**).

**Figure 2:**
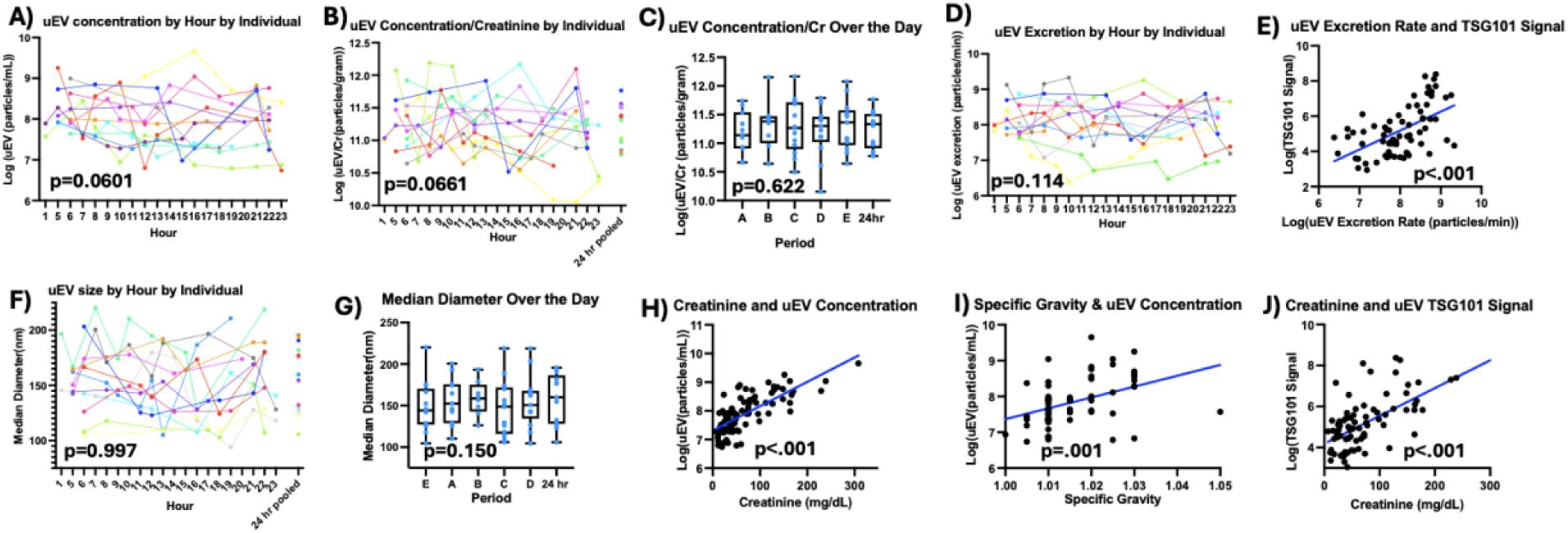
Concentration, Excretion and Size of UEVs. Panel A shows the uEV concentration over the day by individual. Panel B shows the UEV concentration normalized to creatinine by hour each void over the day by individual. Panel C shows boxplots of the UEV concentration normalized to creatinine over the periods (Period A: second morning void. B: All voids between A and 12 pm. C: All voids between 12 pm and 6 pm. D: All voids between 6 pm and 12 pm. E: All voids between 12 am and A). Panel D shows the uEV excretion by hour each void over the day by individual. Panel E shows the trend of UEV excretion rate and UEV TSG101 signal via immunoblotting. Panel F shows the UEV size by hour of each void over the day by individual. Panel G shows boxplots of the uEV median diameter over the periods. Panel H shows the trend of urine creatinine and uEV concentration. Panel I shows the correlation of urine specific gravity and UEV concentration. Panel J shows the correlation of urine creatinine and uEV TSG101 signal via immunoblotting.

### uEV count correlation to normalization markers

Urine is a dynamic biofluid. We determined that the urine creatinine excretion rate is stable in our healthy cohort (**Supplemental Figure 5D**). We then determined the relationships between uEV concentration with urine creatinine concentration, specific gravity, and TSG101 (tumor susceptibility 101), a previously characterized uEV marker. uEV concentrations were positively associated with urine creatinine concentration (**Fig. 2H**) suggesting that uEV concentration in spot urine can be normalized against creatinine concentration in a steady and healthy state. uEV were positively associated with specific gravity tested by a urine dipstick (**Fig. 2I).** The TSG101 immunoblot signal also correlated significantly with urine creatinine concentrations (**Fig. 2J**). Additionally, the TSG101 signal was significantly increased as more uEVs were produced (**Fig. 2E**). The significance for the over the day excretion and relationships between two quantitative variables were calculated via linear mixed modeling (Supplemental Figure 5).

### uEV cargo analysis over a 24-hour period

In order to understand temporal patterns of uEV cargo expression over a 24 hour period, uEV cargo candidates representing different parts of the nephron such as glomerular (CR1, MME, and tubular markers (sodium chloride cotransporter (NCC), aquaporin 2 (AQP2), DPP4, and CD9) (**Fig. 3D)** were selected. NCC, AQP2, and TSG101 were quantified utilizing immunoblotting with an example of Individual B in **Fig. 3A** and all 10 individuals in **Supplemental Figure 6.** NCC signal was normalized to TSG101 signal. NCC/TSG101 signal increased over the course of the day (β = 0.047, p = 0.022, **Supplemental Figure 5**), indicating a modest but statistically significant upward trend across hours after accounting for repeated measures within individuals. When grouping into periods, we found that NCC/TSG101 has a dip at midday and significantly increases thereafter from Period B to C (**Fig. 3B**, p=0.0312). AQP2 does not vary significantly over the course of a day but there is an increase in signal of uEV cargo after noon, but the result is not significant (**Fig. 3C,** p=0.27). Of note, the immunoblotting showed two bands for NCC protein, at the expected sizes for both monomers and dimers (130 and 260 kDa respectively) and both bands were quantified). As mentioned earlier the immunoblot signal of the uEV marker TSG101 correlated significantly with urine creatinine concentrations (**Fig. 2I**).CR1, DPP4, MME, and CD9 were detected with single EV flow cytometry, **see Fig. 3, Supplemental Figure 7 (gating strategy).** The flow panel proteins were also validated via immunoblotting (**Supplemental Figure 8**). No significant temporal patterns were detected over the course of a day for these four nephron markers via flow when grouped into periods (**Fig 3 E-H**) or through linear mixed modeling (**Supplemental Figure 5D**).

**Figure 3:**
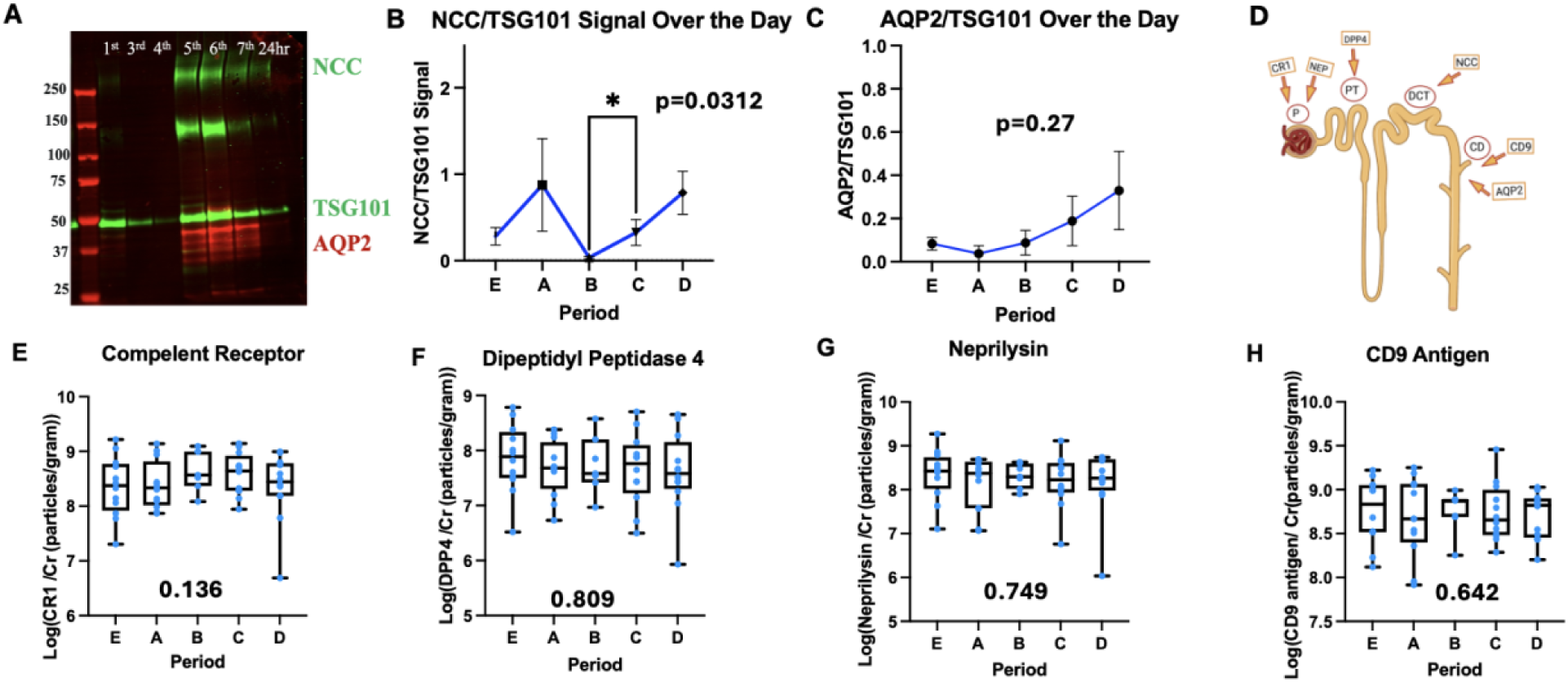
Cargo analysis by immunoblotting and flow cytometry. Panel A shows an example of an immunoblot stained with NCC, AQP2, and TSG101 for individual B’s voids. Panel B shows the NCC normalized to TSG101 signal over the periods. Panel C shows the AQP2 normalized to TSG101 signal over the periods. Panel D shows corresponding to their expression on the nephron For E-F, Single uEV flow cytometry the Cytek Aurora 5-Laser flow cytometer equipped with the Enhanced Small Particle (ESP™) Detection tool. Particle concentration was normalized to urine creatinine values. Panel E shows Complement Receptor 1 over the periods. Panel F shows Dipeptidyl Peptidase 4 over the periods. Panel G shows Neprilysin over the periods. Panel H shows CD9 over the periods. Friedman tests were performed on all.

### uEV counts in the context of food intake over a 24-hour period

Because food and fluid intake impact urine composition, some participants were asked to document their intake using a standardized food log^3^. The relationship between food intake and uEV size and excretion is summarized in a heat map (**Fig. 4A**). Protein diet had a linear trend with uEV size, as did sodium and water intake. Carbohydrate intake had an inverse linear trend with uEV size. uEV excretion had a linear trend with sodium and carbohydrate intake, but not by protein or water intake. We found a trend of increased uEV size with increased fluid intake. As our data was derived from food logs for 9 subjects out of 13 with a free diet, statistical significance was not determined **(Fig. 4B**). Example of a portion of variables from the food log output can be found in **Supplemental Figure 9.**

**Figure 4:**
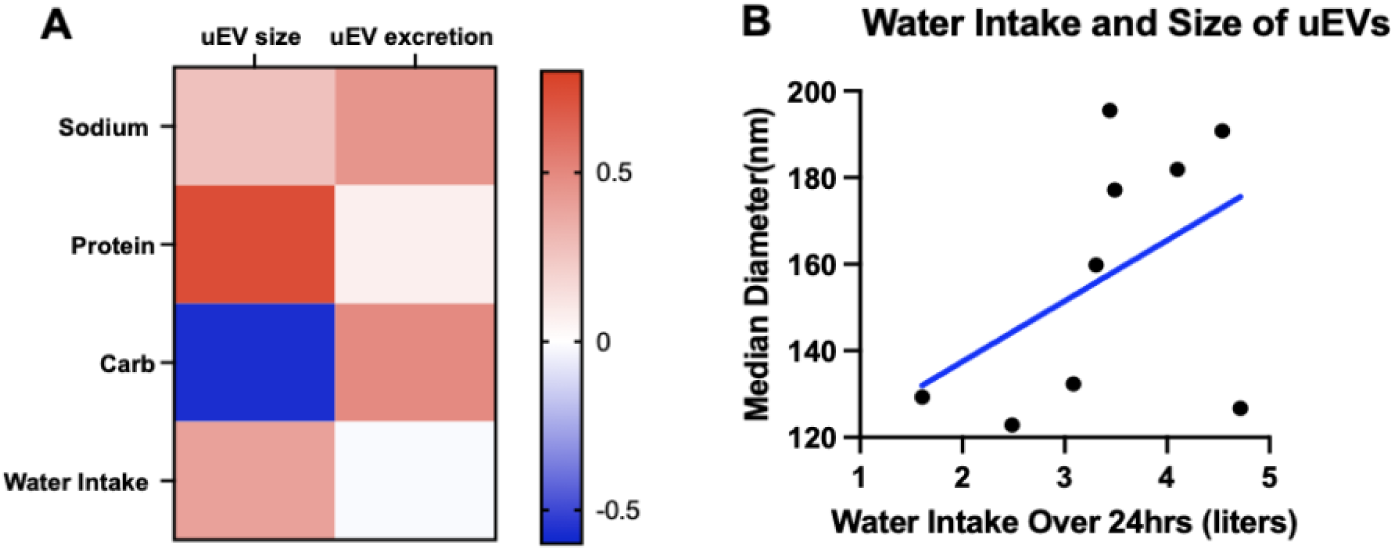
Panel A is a heat map showing the correlation between dietary factors and uEV size and excretion. Correlation values were determined using Spearman’s rank correlation coefficient, which measures the strength and direction of association between two ranked variables. Panel B is a graph showing the association between water intake and uEV size.

## Discussion

Urinary EVs can reflect physiologic and pathologic processes in the kidney and have potential as novel biomarkers of various diseases. Urine as a biofluid carries the benefit of being non-invasive and available in large quantities, making uEVs an appealing target for a liquid biopsy. However, little is known about the overall rate of uEV excretion, which reflects the balance between their secretion and reabsorption in both healthy and disease states. In addition, urine is the most dynamic biofluid with constituents changing day to day and hour to hour. This dynamic process is particularly influenced by diet and fluid intake as well as medications. Normalization strategies for the quantification of urine constituents and uEVs are crucially needed. In addition, temporal patterns of biomarker excretion need to be considered to allow successful translation to the clinic. Since a landmark study published in 1983 in the New England Journal of Medicine (NEJM) demonstrated that the calculated protein-to-creatinine ratio of the second-morning void provided an accurate quantification of daily proteinuria, clinicians have been able to use a spot urine collection to extrapolate kidney health and function across a greater span of time than that during which the single void was produced.^9^ In contrast, not much is currently understood about how well the characteristics of uEVs such as concentration, size, surface markers, and internal contents within a single void reflect the range of those secreted over a longer period of time, e.g. over 24 hours.

This study also employed an optimized uEV separation method. Washes with a low ionic strength buffer were implemented to generate clean uEV pellets to assure that true uEV counts were measured with our Nanoparticle Tracking analysis (NTA) tool and not other non-EV particles and proteins, such as uromodulin. This is supported by other work, which showed that adding uromodulin to urine significantly increased particle counts by NTA^12^. Thus, our uEV separation method increased the robustness of our unique data analysis, allowing accurate assessment of uEV excretion and uEV characteristics.

### uEV concentration, excretion and size variation over the course of a day

Our study is unique as 13 healthy subjects collected each void individually throughout a 24-hour period, along with the exact time collected. To our knowledge only one other study tested timed urine collections for EV analysis in healthy subjects, however participants in that study emptied their bladder at fixed time points, our participants voided spontaneously when participants had the urge to empty their bladder^12^. The uEV collection approach presented here is also in line with the recommendations by the Urine Task Force of ISEV, for example performing a quality control test with a urine dipstick and cooling at proper temperature during urine collection and processing. These recommendations are summarized in the uEV collection card of the “Urine task force” of the International Society for EVs^17^.

As composition and concentrations of urinary constituents vary over the course of the day it is not surprising that uEV concentrations vary over the course of a day as demonstrated in our study. We detected up to 100-fold changes with a decreasing trend close to significance. uEV concentration variation was visualized in individual spaghetti plots visualizing linear mixed modeling accounting for individuals and by sorting data into five prespecified collection periods over the course of the day as done in the landmark paper by Ginsberg et al^9^. uEV excretion is not stable over the day as urine production and time of voiding jointly contribute uEV excretion dynamics A temporal pattern of uEV excretion in timed urine collections of 4 hours over a 24-hour period has previously been described in Sprague-Dawley rats, yet our data set is not directly comparable to the study by Koritzinsky et al. as small uEVs were analyzed utilizing also a different uEV separation method^18^.

Greater urine production is associated with greater uEV excretion in the early morning but less uEV excretion in the evening. This finding is partially in line with the study by Blijdorp et al who demonstrated that the number of uEVs excreted in urine of 11 healthy subjects undergoing a formal water loading test varies inversely with urine excretion^12^. Several in vitro studies suggest proximal to distal tubular cell communication in which uEVs are released and taken up by kidney cells^14^. Depending on the stimulus, here the urine flow rate, changes in uEV release and uptake could be possible.

uEV size was determined by NTA in our cohort and varied between healthy subjects from a median size for an individual void varied from 93.9 to 220.1 nm in diameter but there was no clear trend over the course of the day. In addition, our data is biased towards the detection limits of NTA. However, our results are conforming with other studies who show that majority of uEVs are <200nm^1,15^.

### Consideration for uEV normalization strategies

After blood, urine is the second most studied biofluid for EV biomarker discovery. In contrast to blood samples for which homeostasis is within small margins, urine constituents change significantly over the day^36^. We determined that the urine creatinine excretion is stable over the day across individuals over time (p=0.902). This supports the use of urine creatinine as a normalization marker. In fact, we observed that uEV proportionately increases with higher urine creatinine levels (β = 0.0082, *p* < 0.0001 F), indicating a clear positive trend after accounting for repeated measures within individuals. This finding is supported by other uEV work^12^. Our findings are based on analysis of 91samples from 13 healthy subjects. Blijdorp et al. used cell-free urine from 11 subjects undergoing water loading, but also random urine samples from 14 healthy subjects and 26 patients with polycystic kidney disease (PCKD) a slowly progressive chronic disease ^1^. In all these samples, a correlation between particle counts and creatinine concentration was observed. Of note, our study is the first to study a more pure uEV prep, thus confirming there is a significant positive association between uEV concentration and urine creatinine. To fully understand uEV excretion rates in disease states, other chronic kidney diseases with moderate to large proteinuria and inflammatory disease processes, such as glomerulonephritis, need to be studied. Normalization strategies for acute kidney injury also needs to be explored.

We were also interested in the relationship of uEV concentrations with specific gravity (SG) and TSG101 (tumor susceptibility 101) an uEV marker. We used SG as a surrogate for osmolality measurements and found a positive association between uEV concentration and SG. It is suggested to use osmolarity for normalization for urine smaples^17^. Correlation between particle count and osmolality was reported before ^12^. TSG101 plays a role in the ESCRT pathway, which is involved in the sorting and release of cargo into multivesicular bodies and ultimately into EVs. We saw a positive association between urine creatinine and TSG10. As mentioned earlier Korizinski et al., detected TSG101 excretion patterns for small uEV excretion generated from urine of rodents with ultracentrifugation from timed urine collections of 4 hours ^18^.

The average of uEV concentrations of all individual voids in our collection corresponded to the average result in this “artificial” 24-hour urine collection, supporting that 24-hour urine collection remains the gold standard for correct quantification of uEVs. We conclude that for random urine samples, normalization to creatinine or TSG101 is possible for healthy individuals in which a steady state of glomerular filtration rate exists. If future investigators wish to understand the best time to measure a particular biomarker, a 24 hour urine collection with individual voids as done in our study should be performed.

### uEV protein cargo temporal dynamics over the course of a day

Circadian patterns of extracellular vesicles have been explored previously in various disease pathways and models such as Parkinson’s disease, metabolic dysfunction, obstructive sleep apnea, PCOS, and various cancers have been studied ^19,20,21^. The temporal patterns of extracellular vesicles has also been studied in fluids of various sources including bovine milk, human plasma, stem cells, mouse cell culture, rat urine, and human urine^18,22,23,24,25^. However, few comprehensive studies have been done to characterize the EV protein cargo expression over the day in humans and none before in urine. We detected proteins of glomerular and tubular origin in our sample collection by immunoblotting (NCC, AQP2) and more targeted with single EV flow cytometry (CR1, DPP4, MME, and CD9). Our study found a temporal/diurnal pattern of NCC expression in uEVs with a dip at noon and increase after which was statistically significant. Our findings of a NCC pattern align with a study of five healthy individuals by Castagna et al., who examined the diurnal pattern of NCC and AQP2^22^ using filtration to enrich uEVs. In their study, urine was collected at five fixed time periods rather than from spontaneous voids. In addition, we also observed an afternoon increase in AQP2 similar to this study, but the differences across time points were not statistically significant. Nevertheless, both AQP2 and NCC exhibited similar temporal trends in both studies, supporting some consistency of these patterns^22^.

In a second and targeted analysis of uEV cargo we detected glomerular and tubular protein markers such as CR1, DPP4, MME, and CD9 with single EV flow cytometry in individual voids of our cohort. No significant temporal patterns were detected. Mass spectrometry analysis of individual voids over a 24 hour period would be the next step to provide a more unbiased analysis of uEV protein cargo patterns over a day.

Other groups have looked at diurnal patterns of other EV cargo such as RNA. Circadian rhythm was found to influence adipocyte-derived exosomal miRNAs, which, in turn, regulate insulin sensitivity and metabolic function^24^. Circulating exosomal miRNAs may also serve as indicators of circadian misalignment, potentially playing a significant role in metabolic dysfunction observed in night shift workers^26^. Additionally, the circadian rhythm of adipose-derived stem cells may influence the secretion of small EVs, which modulate inflammation^27^. All these studies support circadian rhythms and further discovery and validation studies are needed.

### uEV excretion and food and fluid intake over the course of a day

Some of our subjects kept a food log for their free diet. Documented fluid intake showed a trend that higher fluid intake corresponds with larger uEVs, but the log was not completed by enough subjects to determine statistical significance. This trend was also observed by Blijdorp et al, which performed a more formal water loading study demonstrating that uEVs’ sizes vary directly with fluid intake and osmolality^12^. The authors concluded that hypotonicity caused the increase in uEV size^12^. Protein consumption of our healthy subjects did not seem to have a difference on uEV excretion; however this needs to be verified with defined diets and also with a larger study to understand the influences of diet on uEV excretion. As high protein diets lead to hyperfiltration, a study needs to be designed particularly to assess how hyperfiltration may influence uEV excretion. Of note, our subjects reported that they did not find the food log burdensome to the 24-hour urine collection process, which is recommended by the guidelines for uEV collection^13^ and we encourage other investigators to include a food log in their studies.

## Limitations

There are limitations in regard to design and technology. Regarding design, we studied a small group of healthy subjects, though our collection represents the largest cohort studied to address uEV diurnal variations and excretion. Our cohort was not large enough to study sex based differences, notable because of previous findings suggesting female subjects have a higher count of urinary EVs compared with male^28^. We added a food log to document free diets of our healthy subjects, but defined diets need to be tested to have enough power to address the effect of food intake on uEV excretion. Kidney mass can also affect uEV excretion^29^, and we cannot rule out an effect of kidney size.

Each EV separation and characterization tool has its technological limitations. We focused on the low centrifugation pellet of uEVs, which includes small and large uEVs^30^. As we did not study an ultracentrifugation pellet, we may have not fully appreciated smaller uEVs. Detection limits of EV tools also need to be considered. NTA has a detection limit for EVs of about 60 nm and is biased towards detecting smaller EVs, depending on standard operating procedure settings. The limit for our small particle flow cytometry tool was about 100-150 nm for EV size, therefore having a bias towards larger uEVs. However, immunoblotting is a bulk analysis tool and should have captured all uEVs in the uEV pellet prepared with our method. We performed targeted uEV protein cargo analysis. A future next step would be to utilize comprehensive tools, such as mass spectrometry, which is an unbiased proteome analysis tool.

In summary, our study is the first and largest physiological study to analyze uEV excretion over a period of 24 hours by collecting each individual void spontaneously in healthy subjects. We observed variations in uEV concentration and found that the association between urine flow and uEV excretion significantly varied across the day uEV protein cargo can reflect diurnal patterns of kidney-specific proteins such as NCC. Larger cohorts in health and disease need to be studied, ideally performing unbiased mass spectrometry and including defined diets. Our study supports use of urine creatinine concentration and TSG101 for normalization strategies for biomarker assessment in cohorts who have a stable kidney function.

