## Supplemental Figure 1 for "Temporal Dynamics of Urinary Extracellular Vesicle Excretion and Cargo in Healthy Subjects over 24 Hours"

Supplemental Figure 1: 11 position table for a Zetaview NTA run. On the far right table there is a removal column where the software gives an error if applicable .

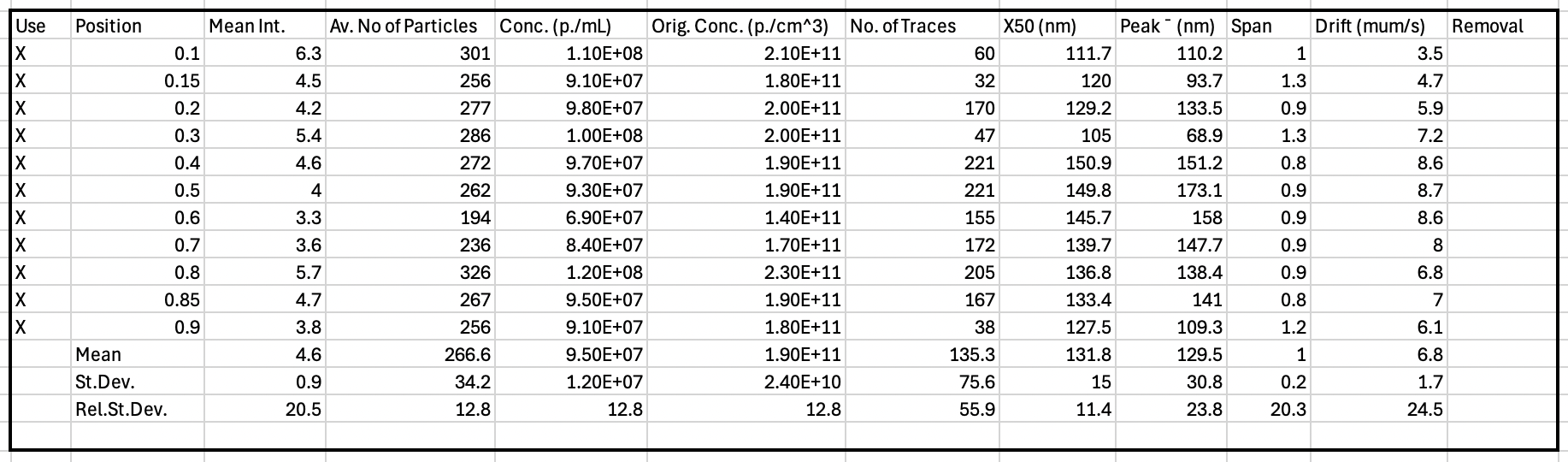
