## Supplementary figures and images for "Temporal Dynamics of Urinary Extracellular Vesicle Excretion and Cargo in Healthy Subjects over 24 Hours"

### Supplemental Figure 2

Supplemental Figure 2: Basic Demographics Table for 13 Healthy subjects
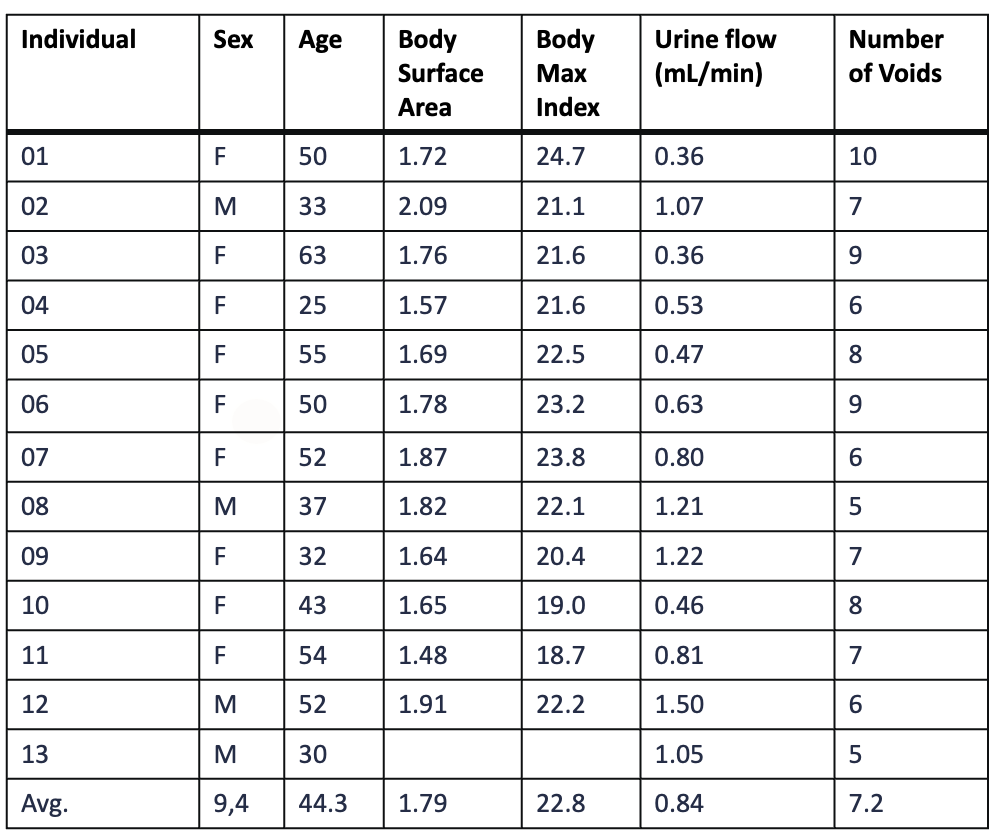

### Supplemental Figure 9

Supplemental Figure 9: Food Log Output for 9 out of the 13 healthy subjects

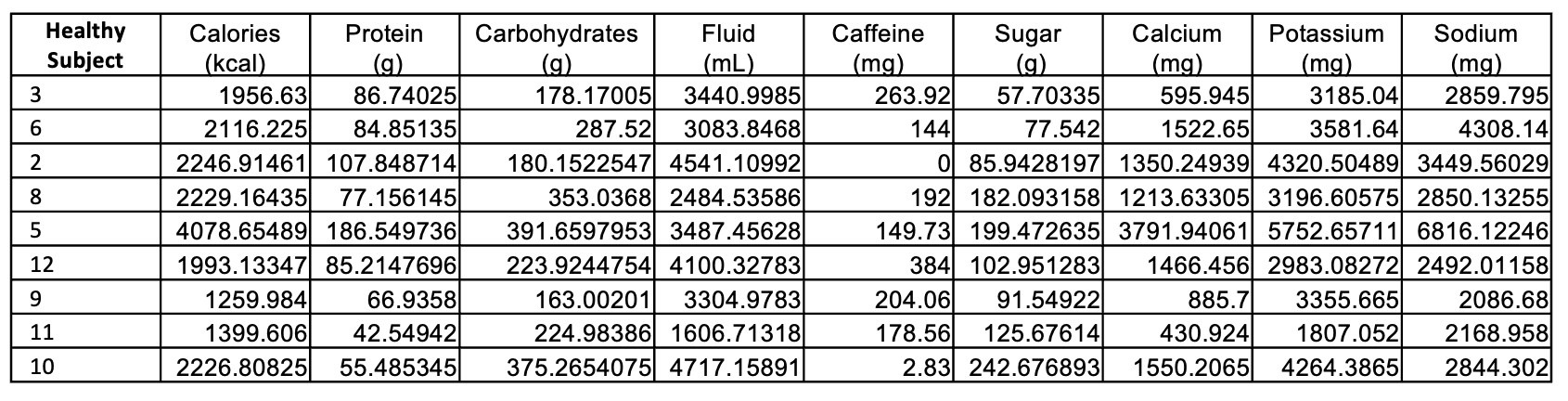
