## Supplemental Figure 3 for "Temporal Dynamics of Urinary Extracellular Vesicle Excretion and Cargo in Healthy Subjects over 24 Hours"

Supplemental Figure 3: Void comparisons for uEV concentration/creatinine (particles/gram) vs 24 hr pooled sample. Nonparametric t test were performed (wilcoxon rank sum).

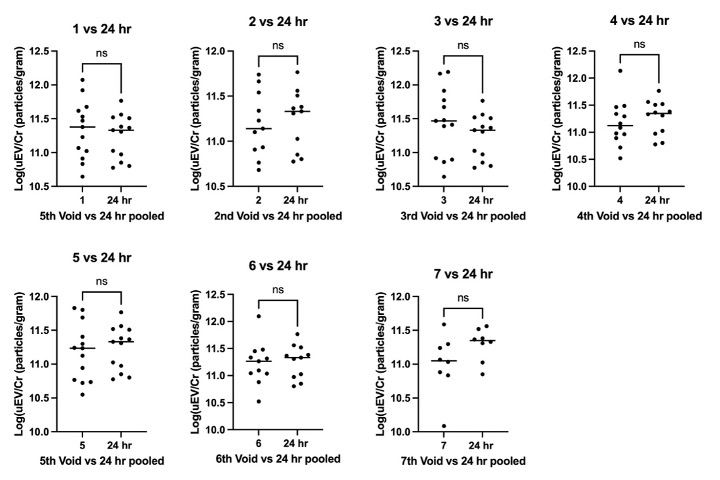
