## Supplemental Figure 4 for "Temporal Dynamics of Urinary Extracellular Vesicle Excretion and Cargo in Healthy Subjects over 24 Hours"

Supplemental Figure 4: Void comparisons for uEV median diameter (nm) vs 24 hr pooled sample. Nonparametric t test were performed (Wilcoxon rank sum).

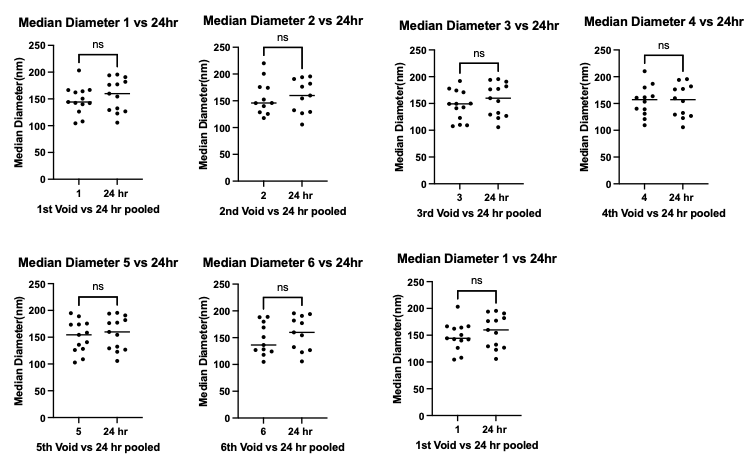
