## Supplemental Figure 5 for "Temporal Dynamics of Urinary Extracellular Vesicle Excretion and Cargo in Healthy Subjects over 24 Hours"

Supplemental Figure 5: Urine production, creatinine concentration, uEV excretion, and linear mixed modeling results. Linear mixed modeling was performed to determine if there was a pattern over the day accounting for the fact that individuals start voiding at different times. The p values of the modeling are reported on the graphs

**B)**

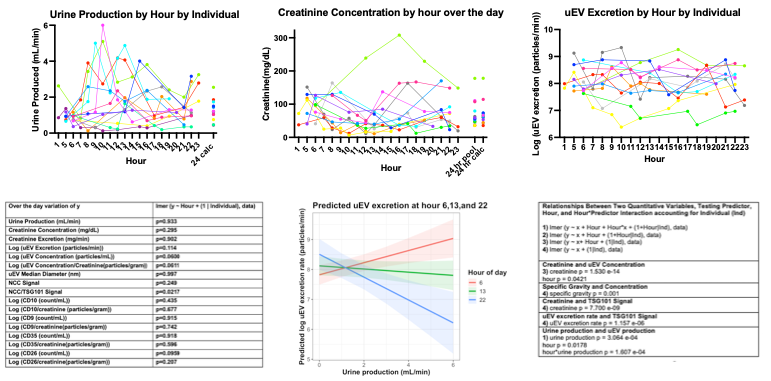

**F)**

**E)**

**D)**

**C)**

**A)**
