## Supplemental Figure 6 for "Temporal Dynamics of Urinary Extracellular Vesicle Excretion and Cargo in Healthy Subjects over 24 Hours"

Supplemental 6: Immunoblots were stained with primary rabbit NCC, anti-rabbit 800 CW, direct conjugate AQP2 680 RD, primary rabbit TSG101, and then anti rabbit 800 CW.

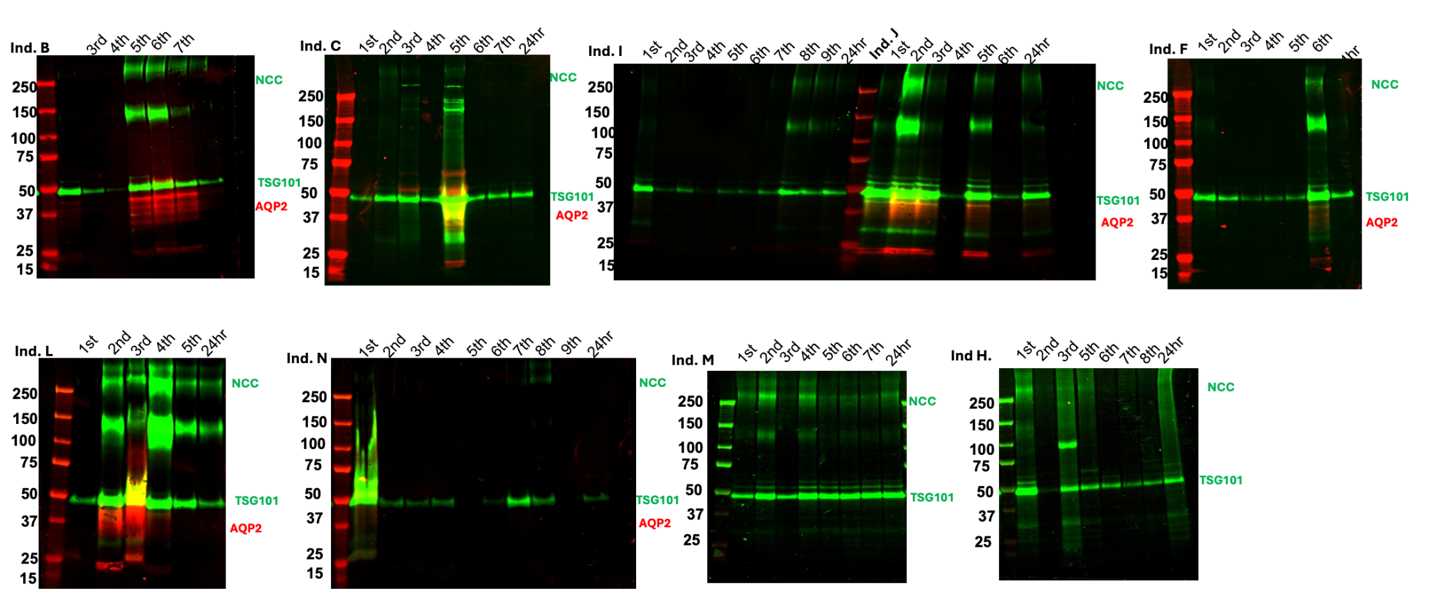
