## Supplemental Figure 7 for "Temporal Dynamics of Urinary Extracellular Vesicle Excretion and Cargo in Healthy Subjects over 24 Hours"

Supplemental Figure 7: Single uEV analysis via flow cytometry equipped with a small particle analyzer. Gating strategy identifies DPP4 positive uEVs, with population percentages identified. (A) Light scatter calibration via FCMPass showing accurate sizing of particles down to <100 nm using a well-fitted scatter-diameter model. (B) Buffer only control, run with each sample batch, demonstrates minimal background fluorescence. (C) Buffer + antibody mix control, run with each sample batch, confirms the absence of background signal and antibody aggregation. (D) Unstained uEV sample demonstrating the presence of uEVs in the sample. (E) Representative stained uEV sample showing DPP4 (conjugated with FITC) fluorescence versus side scatter.

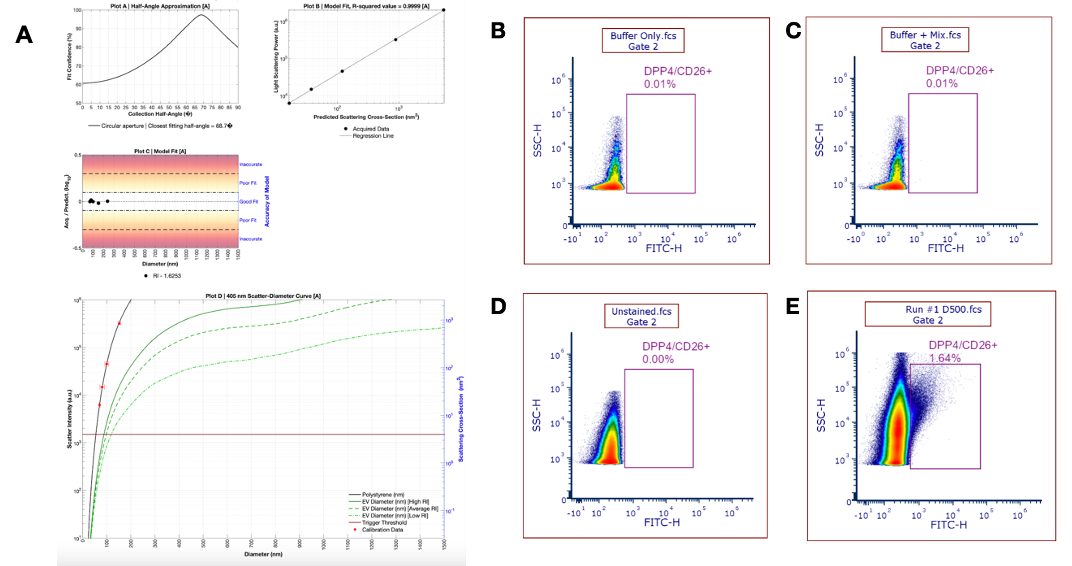
