## Supplemental Figure 8 for "Temporal Dynamics of Urinary Extracellular Vesicle Excretion and Cargo in Healthy Subjects over 24 Hours"

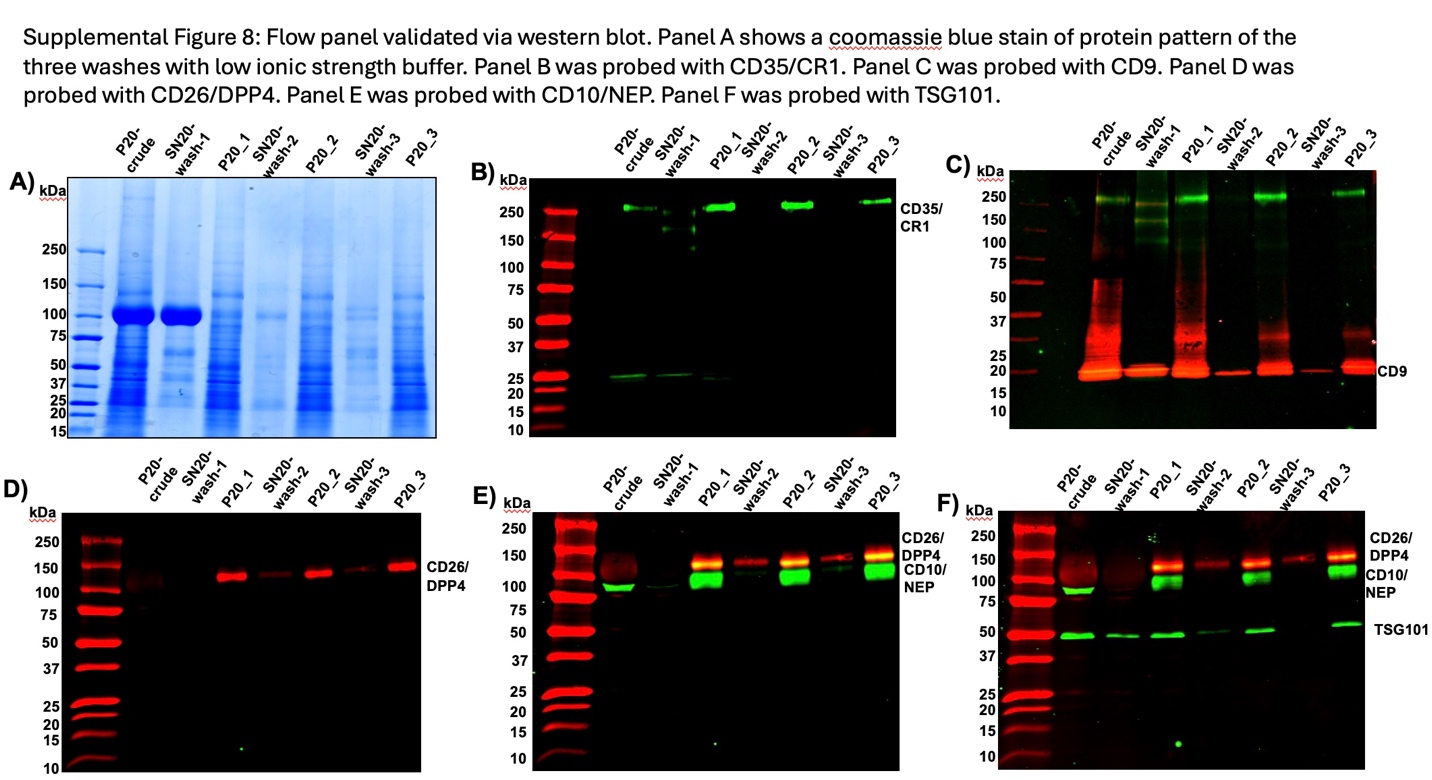

Supplemental Figure 8: Flow panel validated via western blot. Panel A shows a coomassie blue stain of protein pattern of the three washes with low ionic strength buffer. Panel B was probed with CD35/CR1. Panel C was probed with CD9/CR1. Panel D was probed with CD26/DPP4. Panel E was probed with CD10/NEP. Panel F was probed with TSG101

.
